# HuTAge: a Comprehensive Human Tissue- and Cell-specific Ageing Signature Atlas

**DOI:** 10.1101/2024.08.07.606269

**Authors:** Koichi Himori, Zhang Bingyuan, Kazuki Hatta, Yusuke Matsui

**Affiliations:** Institute for Glyco-core Research (iGCORE) Tokai National Higher Education and Research System, Nagoya, Japan; Biomedical and Health Informatics Unit, Department of Integrated Health Science, Nagoya University Graduate School of Medicine, Nagoya, Japan

## Abstract

**Summary:** Ageing is a complex process that involves interorgan and intercellular interactions. To obtain a clear understanding of ageing, cross-tissue single-cell data resources are required. However, a complete resource for humans is not available. To bridge this gap, we developed HuTAge, a comprehensive resource that integrates cross-tissue age-related information from The Genotype-Tissue Expression project with cross-tissue single-cell information from Tabula Sapiens to provide human tissue- and cell-specific ageing molecular information.

**Availability and Implementation:** HuTAge is implemented within an R Shiny application and can be freely accessed at https://igcore.cloud/GerOmics/HuTAge/home. The source code is available at https://github.com/matsui-lab/HuTAge.

## 1 Introduction

Ageing is widely recognized as a major risk factor for a broad range of diseases, including cardiovascular disease, arthritis, neurodegenerative diseases, and cancer (Franceschi *et al*., 2018). To mitigate these adverse effects and enhance health span, a deep understanding of the molecular mechanisms underlying ageing and its involvement in disease is required. The ageing process involves complex molecular and cellular changes, including dysregulated intercellular communication, epigenetic modifications, transcriptional alterations, genomic instability, and defects in telomere maintenance mechanisms, as evidenced by extensive research (López-Otín *et al*., 2023). These processes are characterized by complex interactions among tissues at the macro level and among cells at the micro level. Given these characteristics, understanding the ageing process requires comprehensive data resources with single-cell resolution that span various tissues.

The Tabula Muris Senis (Tabula Muris Consortium, 2020) is a single-cell transcriptomic atlas that spans the entire lifespan of mice and includes data from 23 tissues and organs. This resource is comprehensive and invaluable for examining ageing-related cellular and molecular changes across cell types and tissues. However, comprehensive cross-tissue resources for human ageing research are limited, and a consistent and large-scale single-cell transcriptomic atlas such as the Tabula Muris Senis is lacking. Although studies on ageing phenotypes based on tissue- and cell-specific gene expression in humans exist, these studies often focus on specific cell types within particular tissues (Perez *et al*., 2022; Jeffries *et al*., 2023), which makes systematic com-parisons difficult because of differences in the experimental conditions and assay methods used.

The Genotype-Tissue Expression (GTEx) project (GTEx Consortium *et al*., 2020) provides one of the most comprehensive single datasets with the largest variety of tissue types, encompassing RNA-sequencing (RNA-seq) transcriptome profiles of more than 40 tissues from hundreds of human donors of various ages. Importantly, GTEx has the advantage of including data from multiple tissues from the same individuals. However, while single-cell data are partially included (eight tissues, up to four samples per tissue), they lack sufficient sample size and adequate age-range coverage for ageing research. In contrast, the Tabula Sapiens (Tabula Sapiens Consortium *et al*., 2022) provides cross-tissue single-cell data (24 tissues, up to 13 samples per tissue). Nevertheless, it also has limitations due to the small number of donors and sparse age distribution, making it unsuitable for ageing research when used independently. To address the above limitations, we combined the most comprehensive resources available: GTEx and Tabula Sapiens. One study used the Tabula Muris to deconvolute GTEx data and integrate them both in the context of expression of quantitative trait loci (Donovan *et al*., 2020); however, there have been no studies integrating GTEx and the Tabula Sapiens in the context of ageing. VoyAger (Schneider *et al*., 2024) and GTExVisualizer (Guzz *et al*., 2023) are tissue ageing information resources based on GTEx, but their integration of cellular information is limited.

Here, we introduce HuTAge, a resource that integrates cross-tissue age-related information from GTEx with the cross-tissue single-cell information from Tabula Sapiens. HuTAge aims to (i) provide comprehensive cross-tissue and single-cell transcriptomic data, (ii) facilitate the identification of age-related molecular signatures across various tissues and cell types, and (iii) enable the exploration of age-dependent change s in gene expression, cell-cell interactions, and transcription factor (TF) activities.

## 2 Implementation

HuTAge is a web-based interface developed using the RStudio R Shiny package (https://github.com/rstudio/shiny), designed for exploring and visualizing tissue- and cell-specific ageing signatures. It can be accessed directly from https://igcore.cloud/GerOmics/HuTAge/home and works on any browser. The user guide for the HuTAge is accessible within the application and through Supplementary Materials of the article. The source code and data are available at https://github.com/matsui-lab/HuTAge.

## 3 Features

### 3.1 Overview

HuTAge comprises bulk RNA-seq expression data from 17,382 samples across 30 normal tissues from individuals spanning the age range of 20–70 years obtained from the GTEx project (v.8) and single-cell RNA-seq expression data from 16 normal tissues provided by the Tabula Sapiens project. To allow interactive exploration of the age-dependent single-cell information across human tissues, we constructed four modules: ‘Tissue specificity,’ ‘Cell type composition’, ‘Transcription factor’, and ‘Cell-cell interaction’ (Fig. 1A). The modules are described in detail the following subsections.

**Figure 1.**
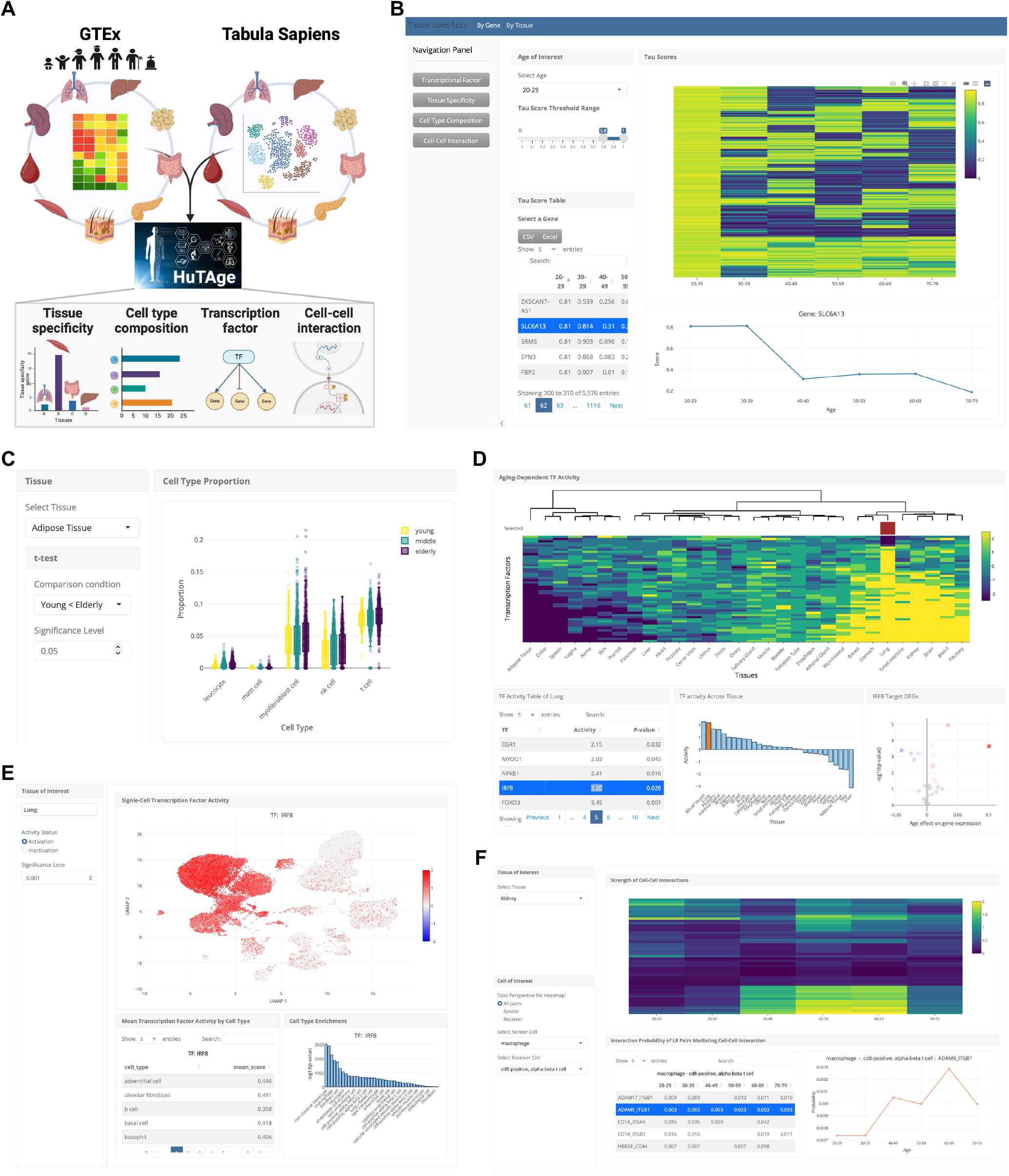
Presentation of HuTAge. (A) Schematic overview of the workflow for exploring tissue- and cell-specific ageing signatures. The web tool HuTAge integrates GTEx bulk organ RNA-seq and Tabula Sapiens single-cell RNA-seq data, providing a comprehensive downstream analysis platform through four distinct modules, as shown below. (B) Heatmap showing tissue-specificity scores of genes across ages in skeletal muscle. Additionally, users can extract genes with high tissue specificity for selected age groups and browse gene information and age-dependent changes in tables and line plots. (C) Box plot displaying age-related changes in cell type composition estimated from GTEx bulk RNA-seq data. (D) Heatmap showing ageing-related changes in TF activity across tissues. (E) Users can map the activity of selected TFs at the single-cell level using feature plots to visually identify which cell types the TFs are enriched in and quantitatively visualize the enrichment in bar plots. (F) Heatmap displaying age-related changes in cell-cell communication strength, with columns representing age groups and rows representing cell and cell pair combinations. Users can also pro-file the activation scores of ligands and receptors mediating these interactions.

### 3.2 Tissue specificity module

The ‘Tissue specificity’ module enables the investigation of age-dependent changes in the tissue specificity of genes (Fig. 1B), which is assessed by calculating the tissue specificity index (tau) for each gene (Yanai *et al*., 2004; Kryuchkova-Mostacci and Robinson-Rechavi, 2017). The ‘By Gene’ tab in HuTAge generates heatmaps that display age-dependent changes in the tau scores of genes, calculated for each age group (Fig. 1B). Users can filter the data by selecting a certain age group and adjusting the tau score threshold, allowing for the extraction of tissue-specific genes for the selected age group. The filtered results can be accessed from a table, and when a gene is selected from the table, a line plot showing the age-dependent changes in its specificity score is provided (Fig. 1B). In the ‘By Tissue’ tab, heatmaps show the tau expression fraction scores (assigned to each tissue and indicating the specificity of the given gene for that tissue) of genes for the selected age group, allowing users to identify in which tissues the genes are specifically expressed.

### 3.3 Cell type composition module

The ‘Cell type composition’ module enables the analysis of age-dependent changes in cell type proportions within tissues of interest (Fig. 1C). Cell type proportions were estimated using Bisque (Jew *et al*., 2020) to deconvolute bulk RNA-seq expression data from different age groups in GTEx, using single-cell RNA-seq expression data from Tabula Sapiens as the reference panel. The ‘Cell type composition’ tab outputs box plots and tables showing cell types with significant differences in proportions between any two of three age groups (young, middle-aged, and elderly) within a single tissue (Fig. 1C). Users can select two age groups for comparison and set the significance level. The ‘Cell marker gene’ tab visualizes age-dependent changes in the expression of cell marker genes in selected tissues and cell types in GTEx data using line plots.

### 3.4 Transcription factor module

The ‘Transcription factor’ module is designed to analyse the age dependency of TF activity across tissues (Fig. 1D, E). TF activity was estimated using the decoupleR method (Badia-I-Mompel *et al*., 2022). The ‘Tissue’ tab visualizes age-dependent changes in TF activity across tissues using heatmaps (Fig. 1D). To achieve this, we conducted an age-dependency analysis using ordinal logistic regression based on the R package ordinal (https://github.com/runehaubo/ordinal). The resulting coefficients were used as inputs to estimate TF activity. Users can extract and visualize TFs that have significant age dependency in tissues of interest, and the extracted TFs can be examined in a table at the bottom of the interface. Upon selecting a TF from this table, a bar plot showing its TF activity distribution across tissues is displayed. Furthermore, target gene expression for the selected TF is presented as a volcano plot, reflecting the coefficients and p-values from the ordinal logistic regression analysis of the target genes. In the ‘Cell’ tab, based on the tissue and TF selected in the ‘Tissue’ tab, the cellular distribution of TF activity is highlighted in a uniform manifold approximation and projection (UMAP) feature plot (Fig. 1E). Specifically, TF activity values estimated for each single cell in the corresponding tissues from Tabula Sapiens are displayed in the UMAP plot. Users can switch between the activation and inactivation states of TF activity in single cells, as well as adjust the significance level to modify the visualization. The table at the bottom of the interface shows averaged TF activity values for each cell type (Fig. 1E). The bar plot integrates p-values from Fisher’s exact test for multiple comparisons, allowing users to identify which cell types are enriched in TF activity (Fig. 1E).

### 3.5 Cell-cell interaction module

The ‘Cell-cell interaction’ module supports the examination of age-dependent changes in cell-cell interaction strength within selected tissues (Fig. 1F). Initially, BulkSignalR was applied to GTEx bulk RNA-seq data to determine ligand-receptor (L-R) pairs that change with age. In brief, we estimated L-R pairs with significant correlations for each age group within the GTEx bulk RNA-seq data. These L-R pairs were input into CellChat (Jin *et al*., 2021) along with single-cell RNA-seq data from Tabula Sapiens to estimate age-specific cell-cell interactions. To extract cells that show age-dependent alterations in cell-cell interaction strength, we calculated communication probability between cell types mediated by the input LR pairs for each age group. The ‘Cell-cell’ tab visualizes the strength of interaction between any cell pair within a selected tissue across different age groups in heatmaps (Fig. 1F). By choosing sender or receiver cells of interest, users can access detailed information about the ligands and receptors mediating these interactions. Upon selecting an L-R pair from the table, a line plot showing the interaction probabilities calculated for each age group is displayed (Fig. 1F).

## 4 Conclusion

We introduced HuTAge, a comprehensive resource that integrates cross-tissue age-related information from GTEx with cross-tissue single-cell information from Tabula Sapiens. By leveraging the strengths of these comprehensive datasets, HuTAge allows multifaceted and comprehensive cross-tissue and cell-type ageing profiling. It identifies targets with varying degrees of association with and specificity to ageing and interprets this information at the cellular level. This integrated resource offers a comprehensive in silico tool for analysing tissue- and cell-specific ageing gene information in humans. Our web-based interface, developed using the RStudio R Shiny package, allows for interactive exploration and visualization of tissue- and cell-specific ageing signatures. Users can access the platform directly from any browser and perform in-depth analyses using the ‘Tissue specificity,’ ‘Cell type composition,’ ‘Transcription factor,’ and ‘Cell-cell interaction’ modules. We plan to further enhance HuTAge by incorporating additional datasets and expanding its analytical capabilities, focusing on improving user experience, integrating more advanced visualization tools, and ensuring that the platform remains a cutting-edge resource for ageing research. HuTAge demonstrates the effective integration of diverse datasets to address the complexities of ageing, providing researchers with a valuable tool to enhance our under-standing of age-related biological processes. This comprehensive and accessible platform aids in the study of gene expression changes across various tissues and cell types, contributing to the advancement of ageing research.

## Data Availability

HuTAge is implemented as a Shiny application and can be accessed at https://igcore.cloud/GerOmics/HuTAge/home. The source code is available at https://github.com/matsui-lab/HuTAge.

