## Supplementary Material for "HuTAge: a Comprehensive Human Tissue- and Cell-specific Ageing Signature Atlas"

**HuTAge user guide**

1. **Navigation panel**

HuTAge consists of four main modules that can be accessed from the navigation panel on the left side of the interface (Fig. S1, box 1).

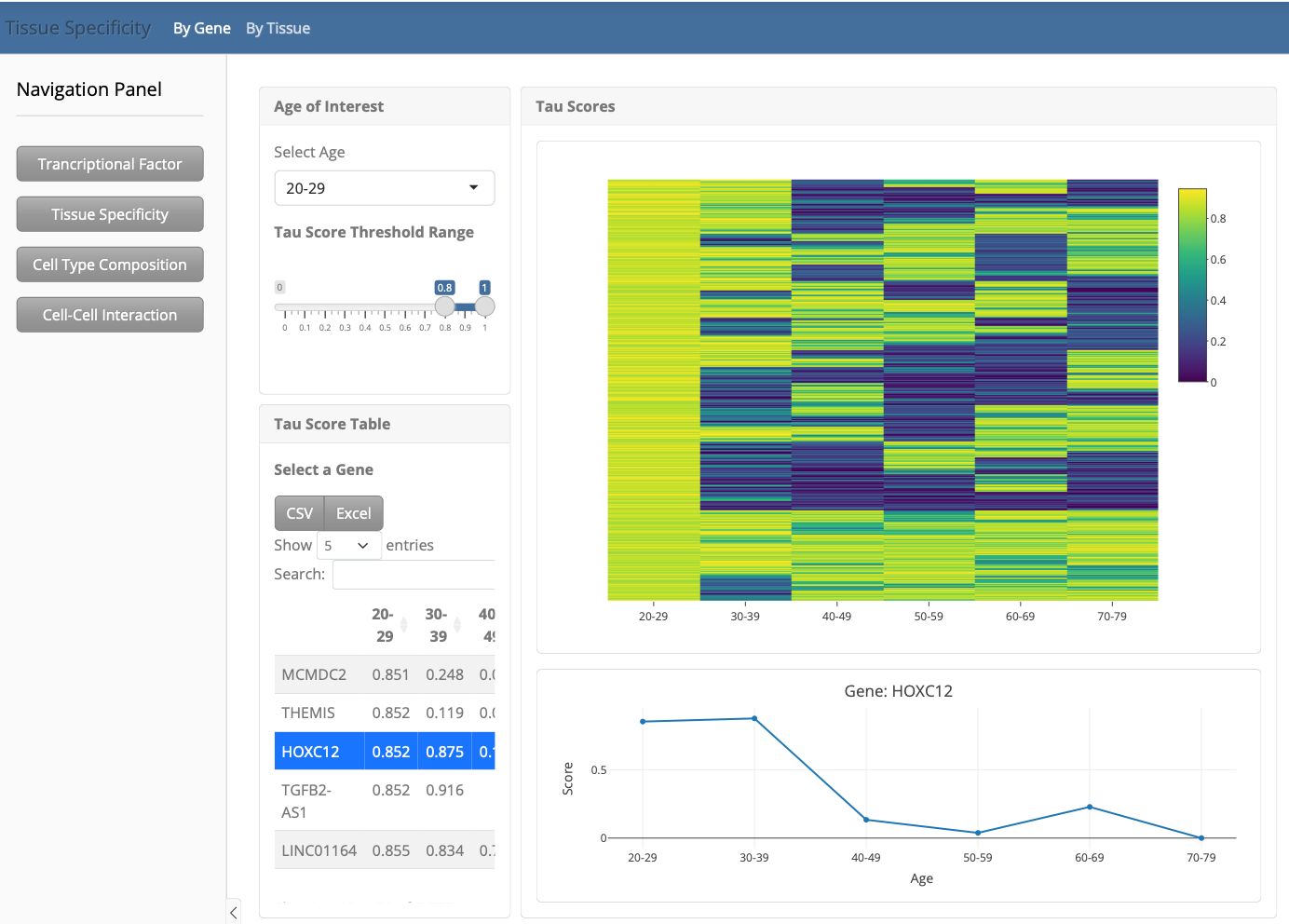

3

1

2

Figure S1. Overview of the HuTAge interface and gene-centric analysis in the tissue specificity module

2. **Tissue specificity module**

The ‘Tissue specificity’ module comprises ‘By gene’ and ‘By tissue’ functionalities that can be switched using the upper tabs (Fig. S1, box 2).

### 2.1. By gene tab

The ‘By gene’ tab displays a heatmap showing the tau scores of genes (rows) across different age groups (columns), accompanied by a corresponding table (Fig. S1). When a gene is selected from the table (Fig. S1, box 3), a line plot representing the selected gene is displayed (Fig. S1).

At the top of the sidebar, the user can select the age and adjust the tau score threshold (Fig. S2, box 1), after which all plots and tables are updated accordingly (Fig. S2). In the given example, only the heatmap and table for genes with a tau score of > 0.9 in the 50 - 59 age group are displayed.

1

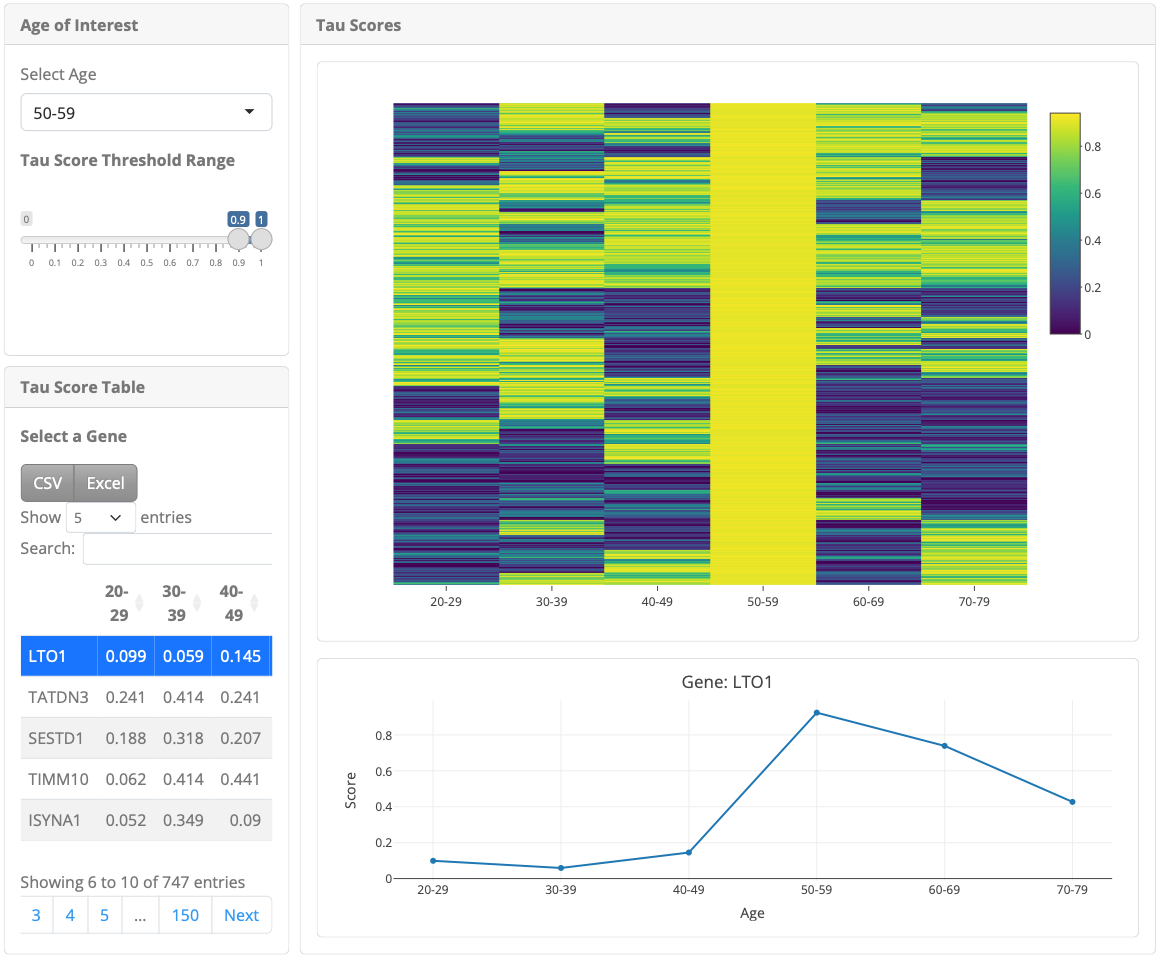

Figure S2. Gene-centric analysis in the tissue specificity module

### 2.2. By tissue tab

In the ‘By tissue’ tab, a heatmap displays the tau expression fraction scores of genes (rows) across tissues (columns) for **the age group selected in the ‘By gene’ tab** (Fig. S3, box1). A table corresponding to the heatmap data (Fig. S3) is also shown. When a gene is selected from the table (Fig. S3, box 2), a line plot depicting its expression across tissues is displayed (Fig. S3).

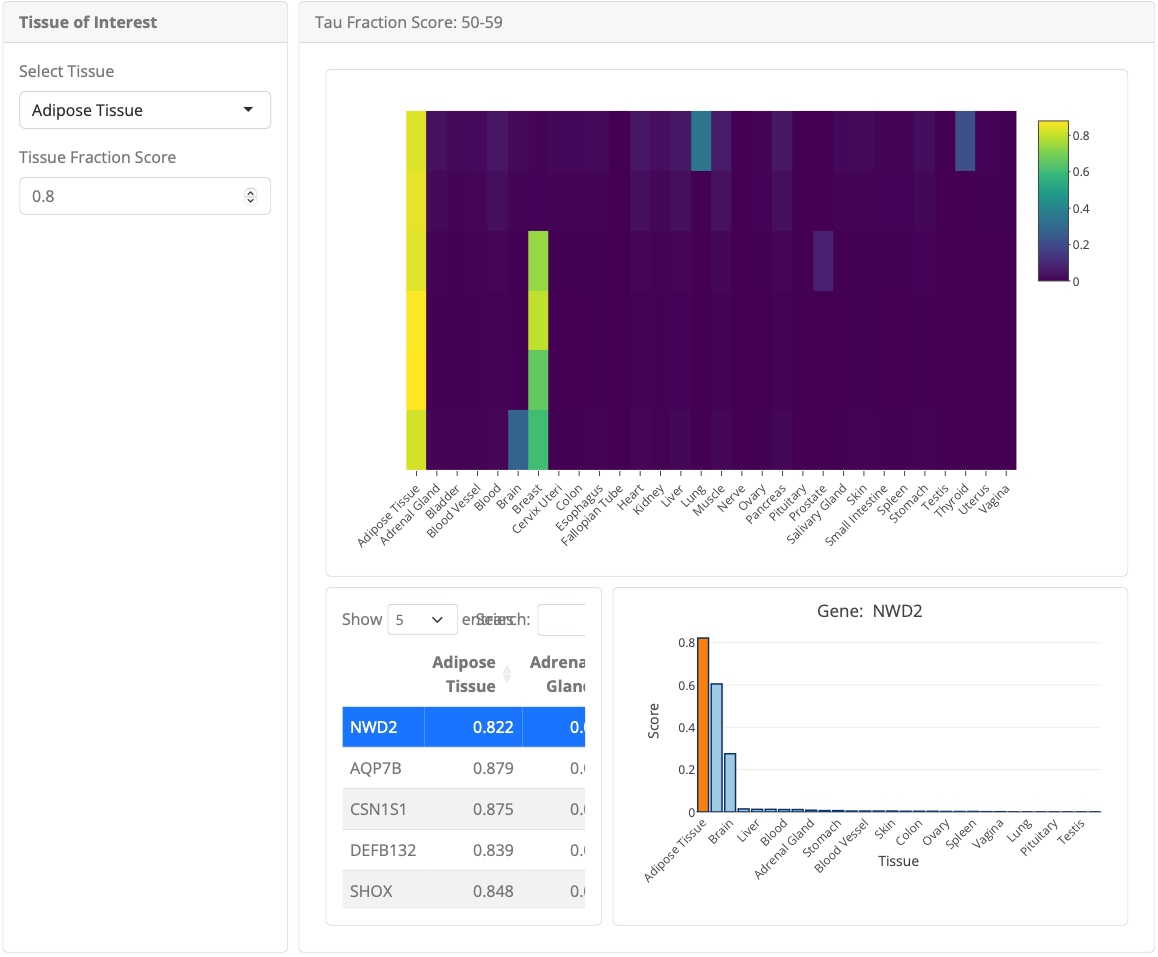
Figure S3. Tissue-centric analysis in the tissue specificity module

1

2

The user can select a tissue from the sidebar and adjust the tau expression fraction score threshold (Fig. S4, box 1), after which all plots and tables are updated accordingly (Fig. S4). In the given example, a heatmap and table showing only the genes with a tau expression fraction score of ≥0.8 in the brain are displayed (Fig. S4).

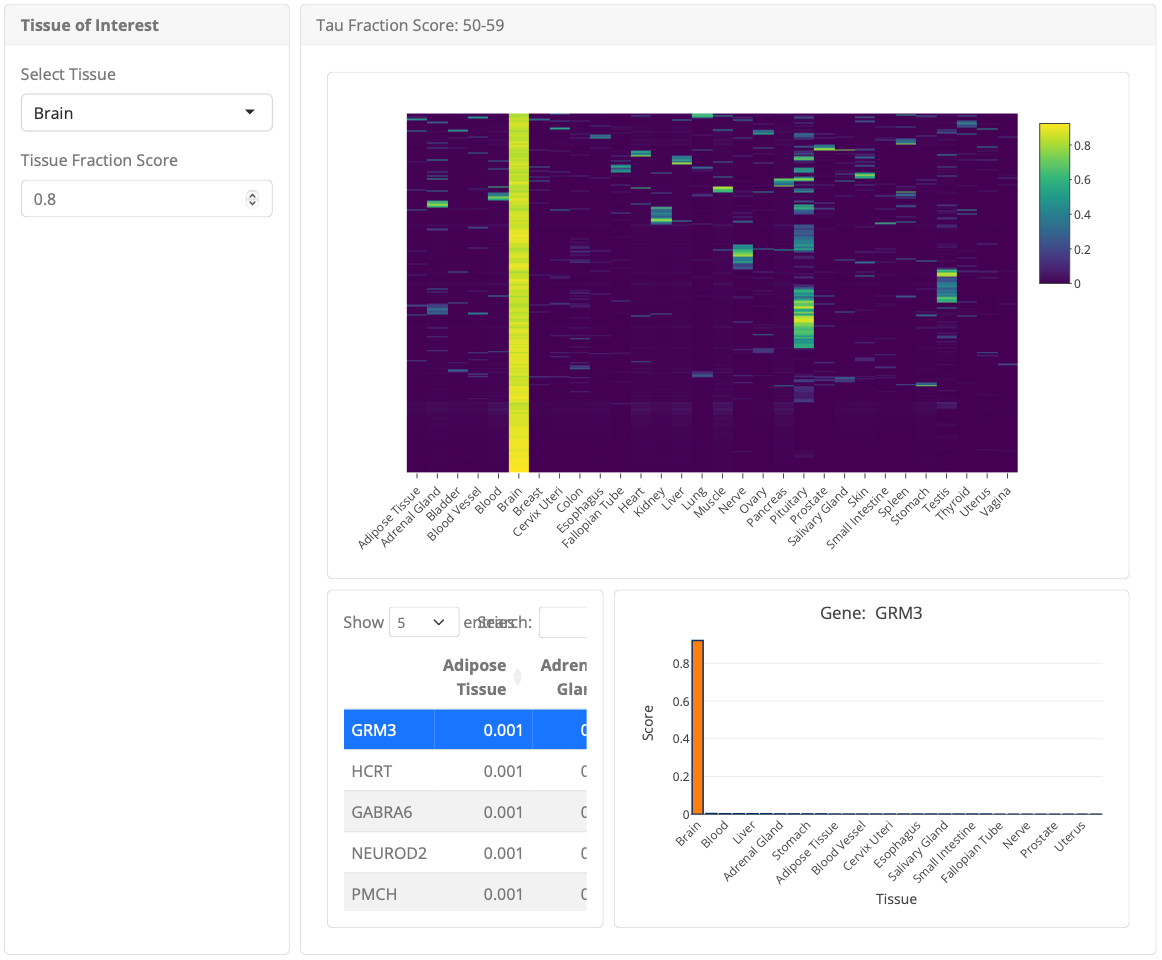
Figure S4. Adjusting tau score threshold for tissue-centric analysis in the tissue specificity module

1

3. **Cell composition module**

The ‘Cell composition’ module comprises two tabs: the ‘Cell type proportion’ tab and the ‘Cell marker gene’ tab, which can be switched.

### 3.1. Cell type proportion tab

In the ‘Cell type proportion’ tab, box plots display the proportions of cell types that show significant changes in gene expression across different age groups (Fig. S5). Corresponding tables show the mean values for these proportions (Fig. S5).

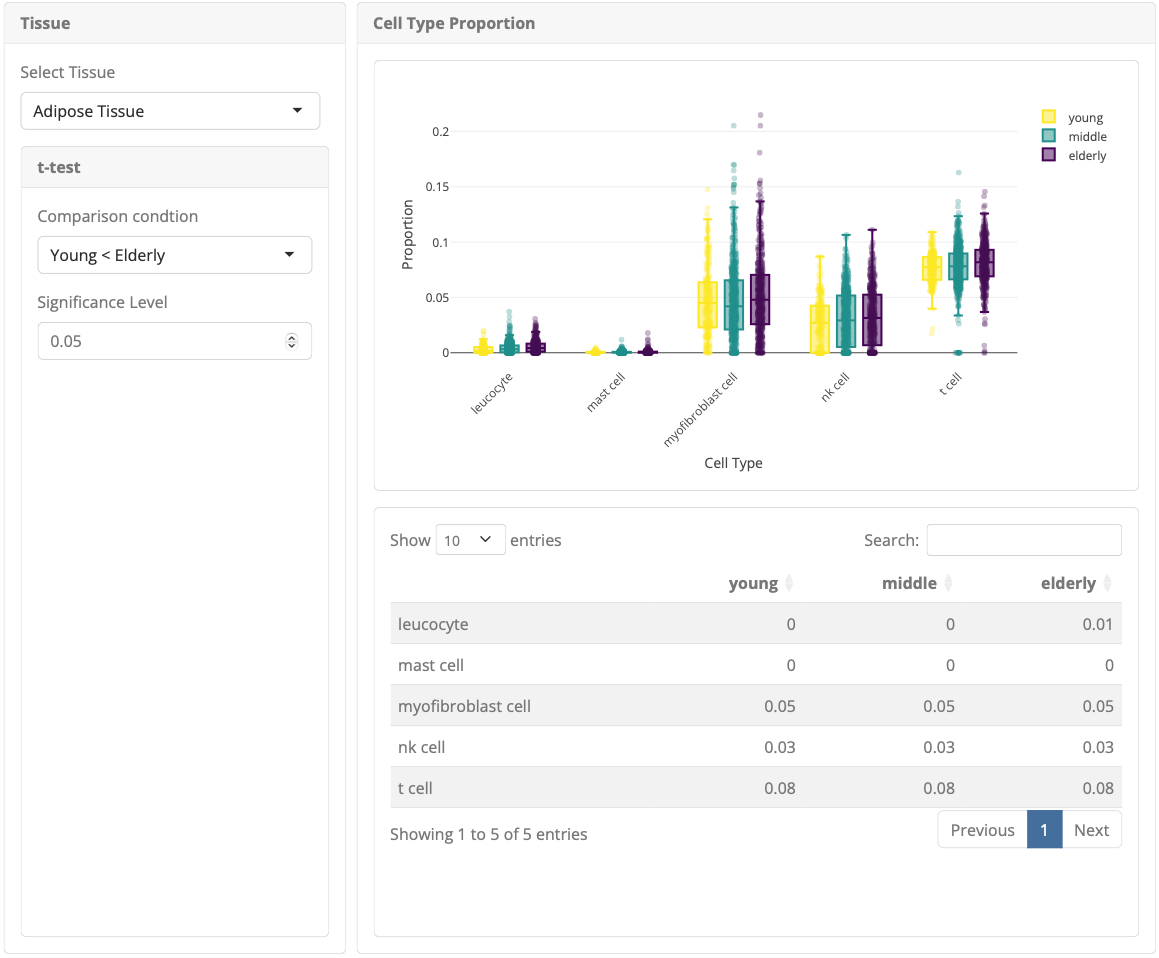
Figure S5. Cell type proportion analysis in the cell composition module

The user can change the tissue in the sidebar (Fig. S6, box 1). By selecting the comparison groups and direction (Fig. S6, box 2), **only those cell types with significant differences in proportions in the chosen direction between the two groups selected will be displayed** (Fig. S6). Additionally, the user can adjust the significance level (Fig. S6, box 2). As an example, Figure S5 shows cell types with significantly increased proportions in the ‘Elderly’ group compared to the ‘Young’ group. Changing the selection to ‘Young > Middle’ (Fig. S6) will display cell types with significantly increased proportions in the ‘Young’ group compared to the ‘Middle-aged’ group.

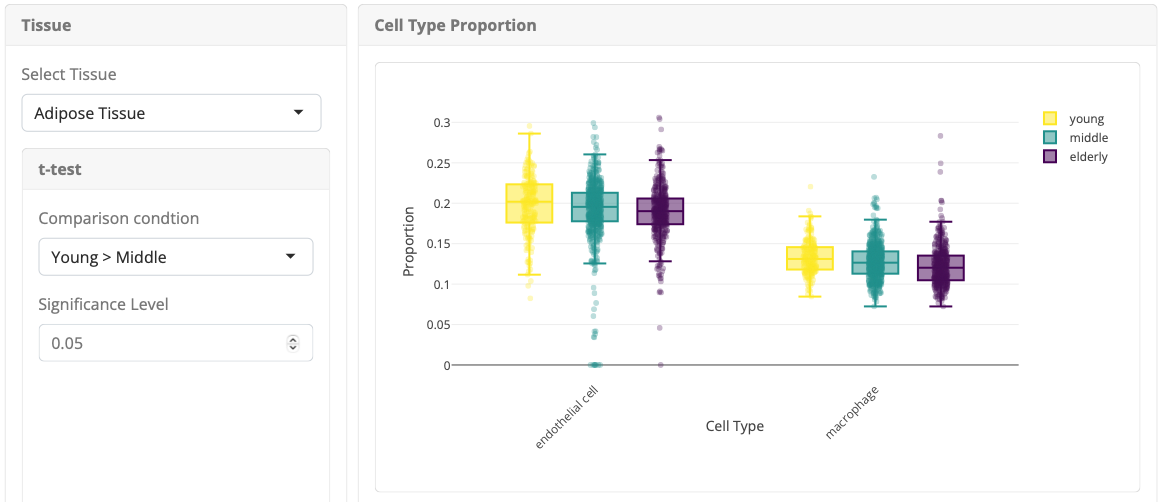
Figure S6. Sidebar Controls for cell type proportion analysis in the cell composition module

2

1

### 3.2. Cell marker tab

In the ‘Cell marker gene’ tab, a line plot (Fig. S7) displays the gene expression values of marker genes for the selected tissue-cell type in corresponding bulk tissues from GTEx, categorized by age group. When a specific marker gene is selected, a box plot shows the gene expression values of single samples grouped by age (Fig. S7).

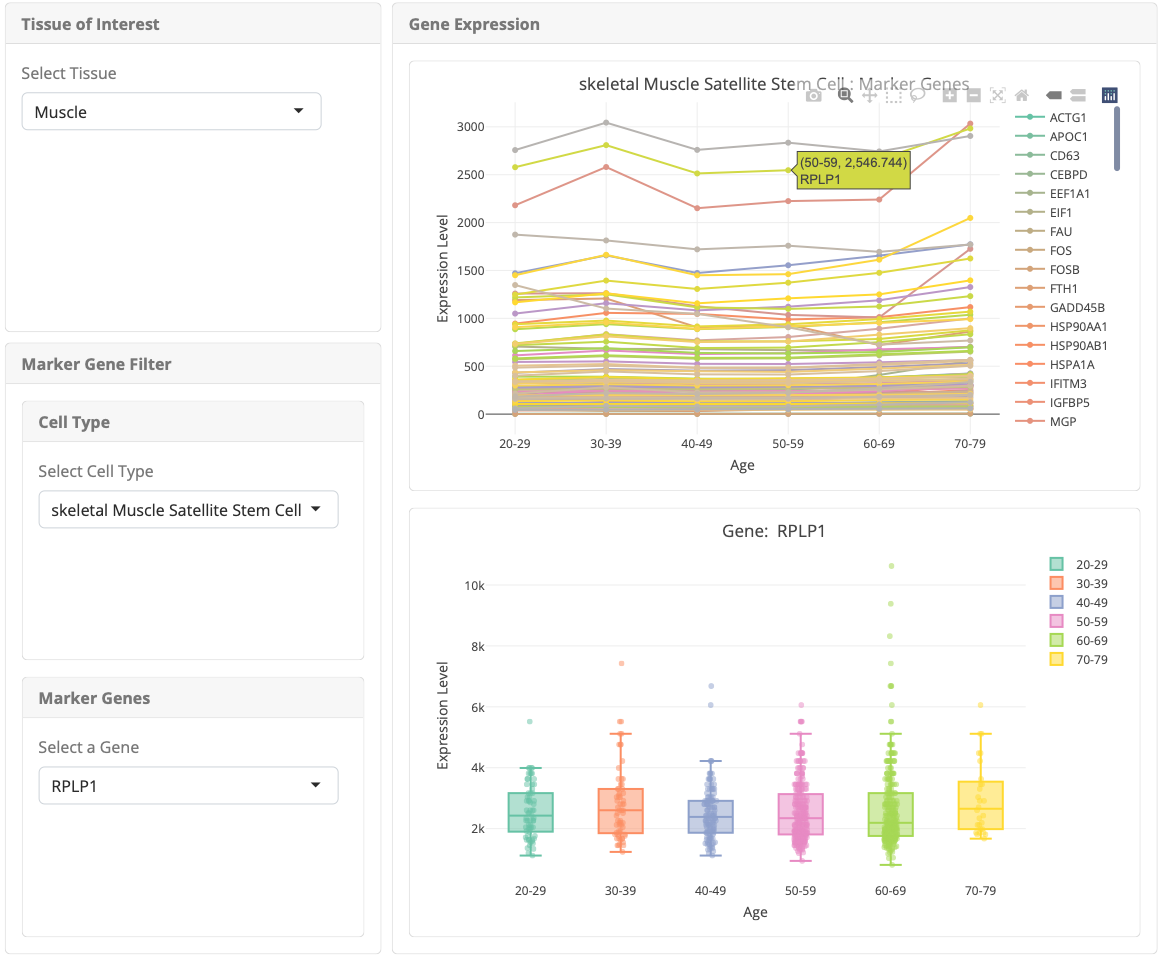
Figure S7. Cell marker gene expression analysis in the cell composition module

4. **Transcription factor module**

The ‘Transcription factor’ module consists of two tabs: the ‘Tissue’ tab and the ‘Cell’ tab, which can be toggled at the top.

### 4.1. Tissue tab

The ‘Tissue’ tab displays a heatmap (Fig. S8) showing transcription factors (rows) and tissues (columns) with age-dependent changes in activity. The colour intensity of the heatmap reflects the degree of age-dependent changes in transcription factor activity, with higher values indicating increased activation and lower values indicating inactivation in response to ageing. A corresponding table (Fig. S8) provides detailed data for the selected tissue. After selecting a transcription factor from the table (Fig. S8, box 1), its data across all tissues are displayed in a bar plot (Fig. S8). Additionally, a volcano plot shows the age-dependency statistics (coefficients and –log10 *p*-values) of the target genes for the selected transcription factor in the corresponding GTEx tissue.

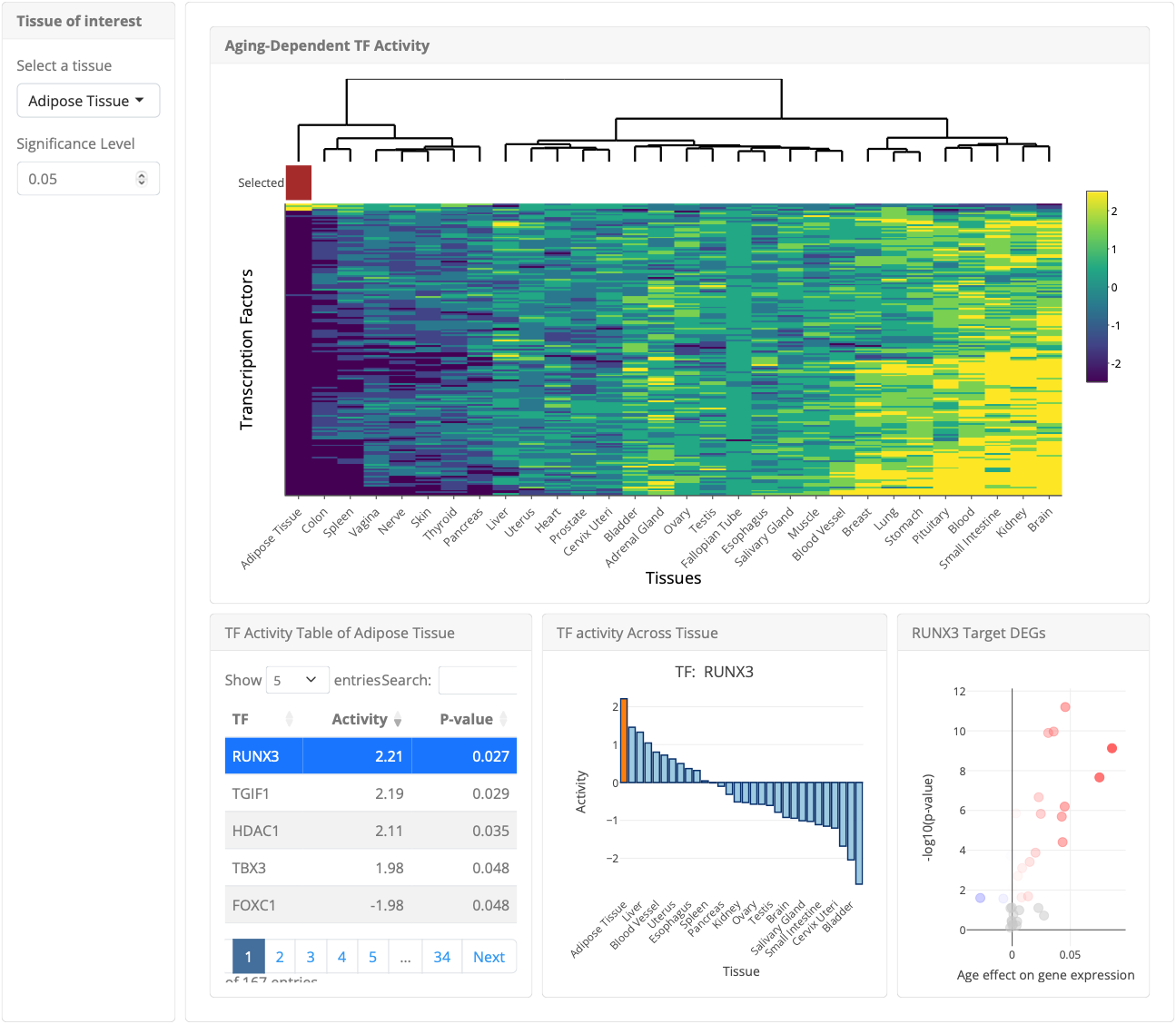
Figure S8. Tissue-based transcription factor analysis in the transcription factor module

1

### 4.2. Cell tab

The ‘Cell’ tab displays a feature plot showing the cellular distribution of the transcription factor selected in the ‘Tissue’ tab and a table showing the average transcription factor activity in each cell type (Fig. S9). The bar plot quantifies the cellular enrichment of transcription factor activity according to a Fisher’s exact test.

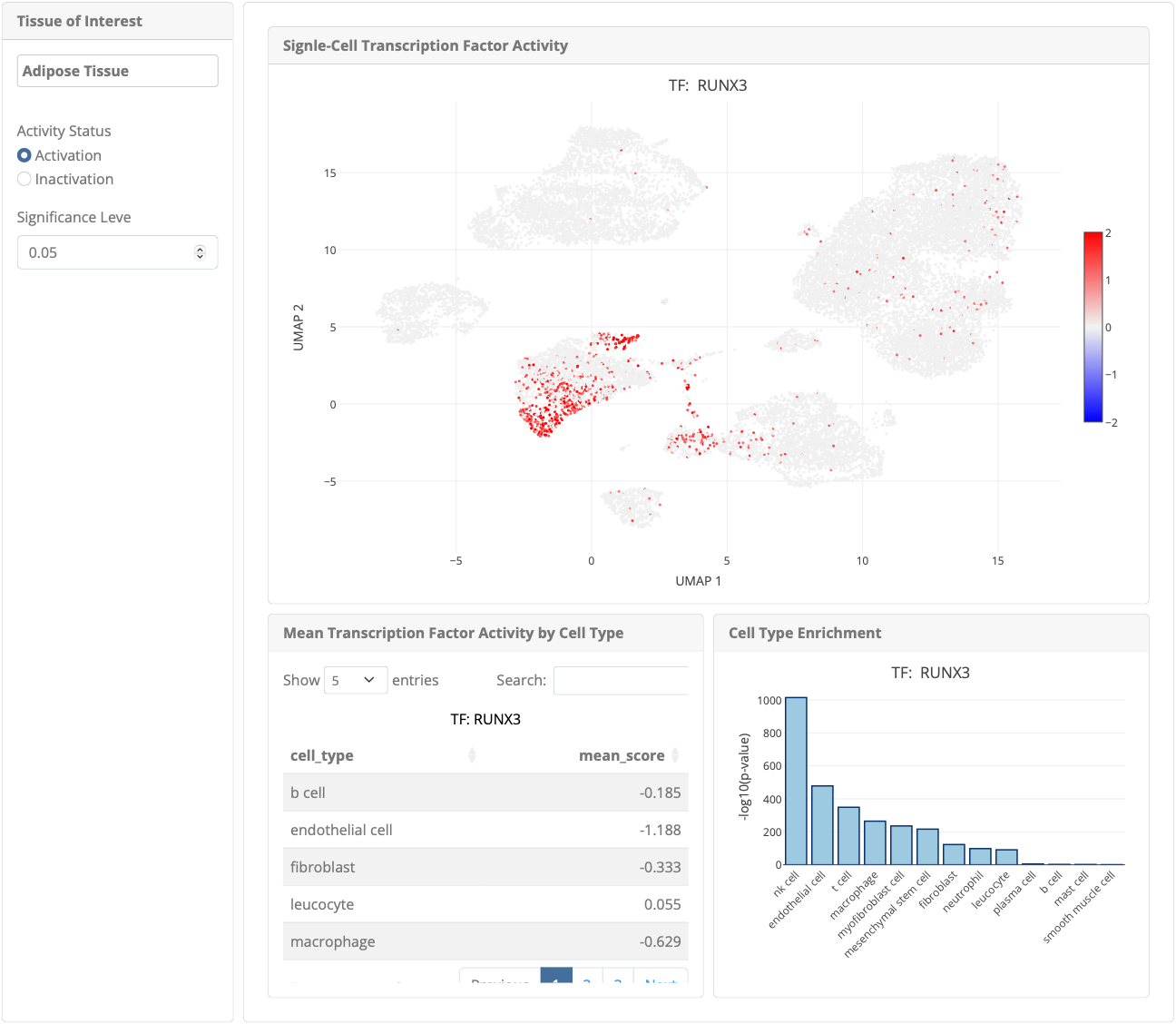
Figure S9. Cell-based transcription factor analysis in the transcription factor module

At the sidebar, the user can switch between cells with activated and inactivated transcription factor activity using ‘Activity Status’ selection buttons (Fig. S10, box 1).

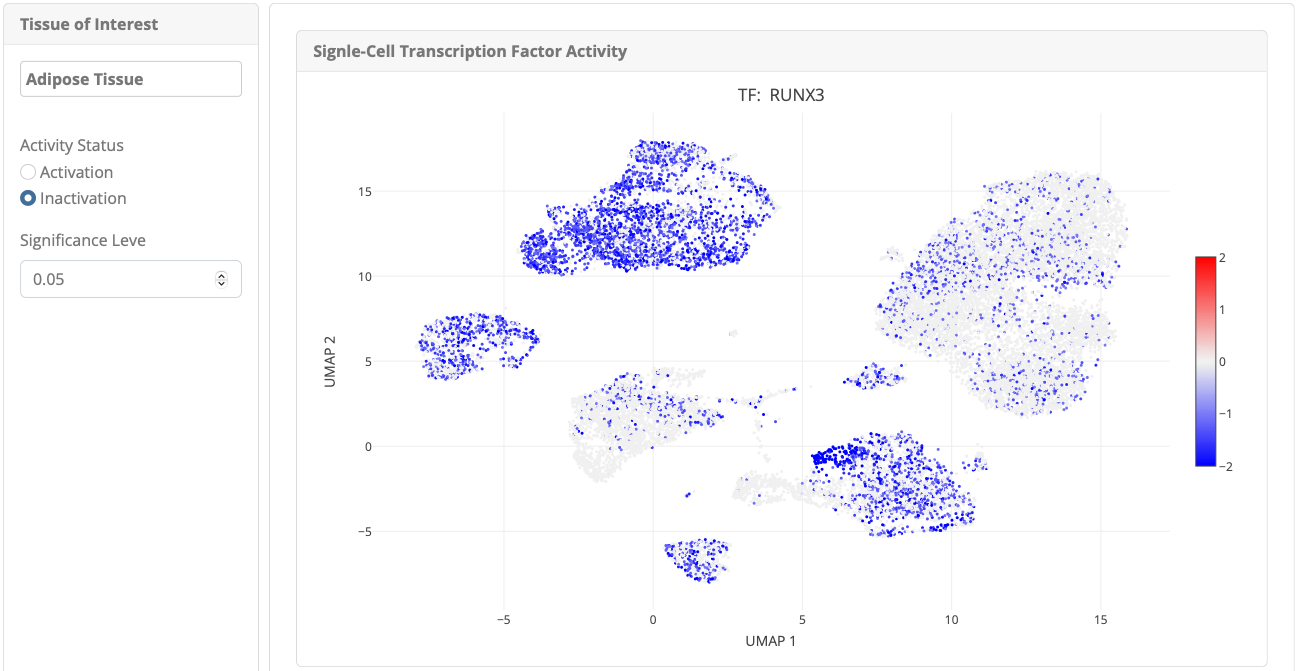
Figure S10. Sidebar Controls for cell-based transcription factor analysis in the transcription factor module

1

5. **Cell-cell interaction module**

The ‘Cell-cell interaction’ module displays a heatmap showing age-dependent changes in the strength of cell-cell interactions among all cell pairs (rows) within a selected tissue across different age groups (columns) (Fig. S11).

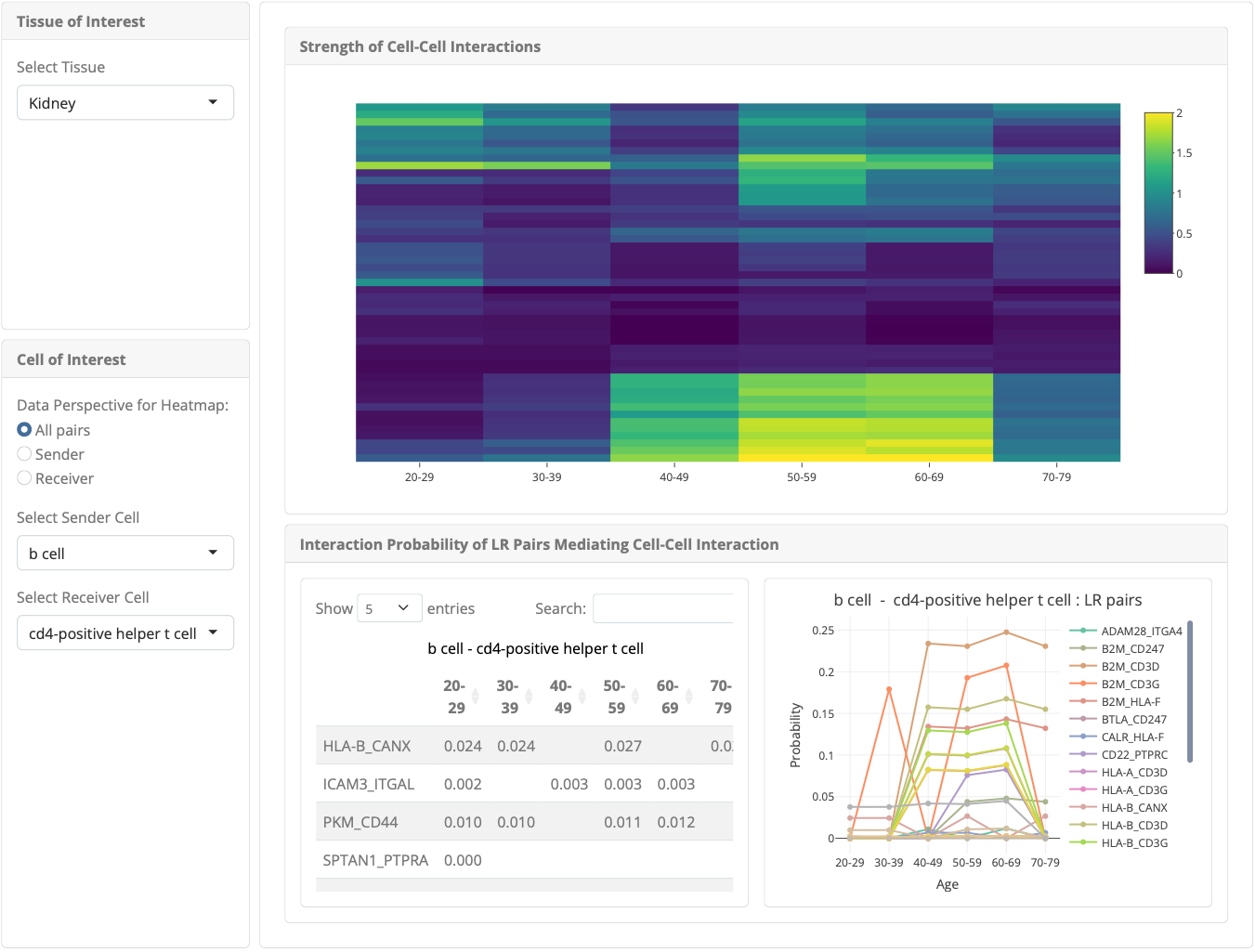
Figure S11. Cell-cell interaction module

Selecting ‘Sender’ radio button in the sidebar (Fig. S12, box 1) will result in a heatmap of all pairs involving the sender cell selected below and other cells, whereas selecting ‘Receiver’ will retrieve a heatmap of pairs involving the receiver cell selected below and other cells. In the given example, the heatmap shows all pairs of cell types in the muscle (Fig. S11). If ‘Receiver’ is selected (Fig. S12, box 2), the heatmap will display pairs where all cells are senders, and cd8-positive cells are receivers (Fig. S12). When one sender cell and one receiver cell are selected, a table presenting the probability of ligand-receptor pairs mediating these interactions by age group is generated (Fig. S12). Selecting a ligand-receptor pair from the table will display a corresponding line plot (Fig. S12).

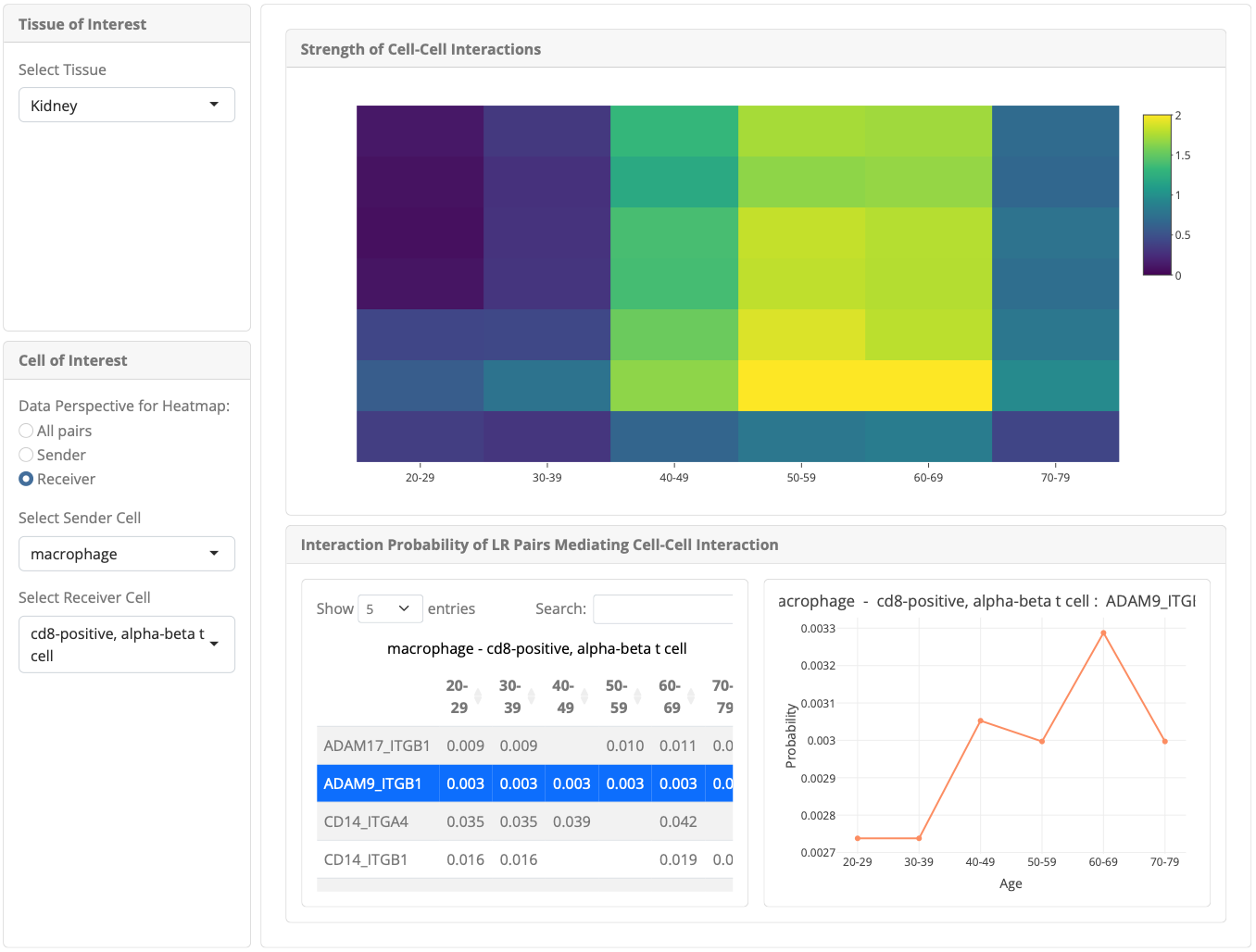
Figure S12. Sidebar controls in the cell-cell interaction module

2

1
